# Arctic charr phenotypic responses to abrupt temperature change: an insight into how cold water fish could respond to extreme climatic events

**DOI:** 10.1101/2020.07.01.182501

**Authors:** Oliver E. Hooker, Colin E. Adams, Louise Chavarie

## Abstract

Phenotypic plasticity, the ability of an organism to express multiple phenotypes in response to the prevailing environmental conditions without genetic change, may occur as a response to anthropogenic environmental change. Arguably, the most significant future anthropogenic environment change is contemporary climate change. Given that increasing climate variability is predicted to pose a greater risk than directional climate change, we tested the effect of a water temperature differential of 4 ºC on the Arctic charr phenotypic response within a generation. We demonstrate that Arctic charr phenotype can respond rapidly and markedly to an environmental cue. The plastic response to different temperature regimes comprised a shift in the mean phenotype coupled with a reduction in the between-individual phenotypic variation in the expressed head shape. The magnitude of shape difference was cumulative over time but the rate of divergence diminished as fish became larger. Individuals raised in the elevated temperature treatment expressed a phenotype analogous to a benthivorous ecotype of this species rather than that of the parental pelagic feeding form. The response of cold-water freshwater species to temperature change is likely to be an interaction between the capacity of the organism for phenotypic plasticity, the speed of mean change in the environment (e.g., temperature), and the degree of short interval variation in the environment.

## INTRODUCTION

In nature, the expression of intraspecific, discrete, alternative phenotypes can result from modulation by the environment through plasticity (Adams, Woltering & Alexander, 2003) or from the emergence of evolutionary groups with diverging gene pools (Wu & Ting, 2004), or a combination of both (Nosil, 2012). Phenotypic plasticity is the ability of a single genotype to express multiple alternative phenotypes in response to different environmental conditions (Pigliucci & Preston, 2004). Plasticity itself is a trait, present across a broad range of taxa (Lubchenco & Cubit, 1980; Corno & Jürgens, 2006; Berg & Ellers, 2010). It can be instantaneous, anticipatory or delayed, permanent or reversible, adaptive or non-adaptive, beneficial or harmful, passive, discrete, continuous, and generational (Whitman & Agrawal, 2009). It is known to facilitate the expression of novel phenotypic traits (Skúlason *et al*., 2019) upon which selection may then act (Ghalambor *et al*., 2007). Thus, phenotypic plasticity has an important role in the evolutionary processes of organisms that exhibit plasticity because variation fuels evolution (Schoener, 2011).

Identifying pathways along which phenotypic plasticity can arise is still a major topic for inquiry in evolutionary ecology, since the patterns and processes that underlie variation are multifaceted and highly variable. Despite the complex nature of phenotypic plasticity, there are some abiotic factors that are known to be ubiquitous in driving variation in ecosystems, one of which is temperature (Mcphee, Noakes & Allendorf, 2012; Noble, Stenhouse & Schwanz, 2018). Many plants and animals have the capacity for a plastic response to temperature, as it is unlikely that rapid environmental change would induce an equally rapid genetic response (Orizaola & Laurila, 2009; O’Dea *et al*., 2016; Merilä & Hendry, 2014). A rapid change in temperature could induce greater levels of phenotypic variation among individuals within a population by exposing previously hidden (cryptic) genetic variation or by inducing new epigenetic changes (O’Dea *et al*., 2016). Accordingly, the response of an organism to temperature can include variation in the mean expressed phenotype but it can also include change in the variance of phenotypes expressed in a population (O’Dea *et al*., 2019).

As ectotherms, fish are particularly sensitive to change in temperature (Neuheimer *et al*., 2011). A phenotypic trait in fish that is strongly linked with temperature is growth (van der Have & de Jong, 1996; Kingsolver, Izem & Ragland, 2004). However, the subsequent effect of temperature and induced growth rate heterogeneity (e.g., early developmental and juvenile growth) on the modulation of morphology is not a mechanism fully understood, despite being observed to affect morphology modulation in a number of species (Olsson, Svanbäck & Eklöv, 2006; Jacobson, Grant & Peres-Neto, 2015; Heino, 2014). A few studies have looked at the potential for temperature change to result in heterochrony in the developmental and ontogenetic processes that modulate the expression of functional traits (Parmesan, 2006; Charmantier *et al*., 2008), whilst others have attempted to predict the morphological outcome of temperature induced plasticity (Mcphee, Noakes & Allendorf, 2012). However, the ecological consequences that result from temperature induced phenotypic change has received little attention ectotherms at higher latitudes (Ramler *et al*., 2014; Burggren, 2018). As the expression of some discrete functional phenotypes in some species is plastic, and by definition modulated by variation in the natural environment, it is presumed that they will also respond to anthropogenic modifications of that environment. The most obvious and far reaching anthropogenic modification of the contemporary environment is that of climate change. The northern regions are warming at a rate twice the rest of the planet and given that a rise in water temperature is predicted to occur, cold-water freshwater species will be foremost affected by climate change (Solomon, 2007; Poesch *et al*., 2016; Heino, Virkkala & Toivonen, 2009).

The Arctic charr (*Salvelinus alpinus*) has a circumpolar distribution, is the most northern species of freshwater fish, and displays highly variable phenotypes (Klemetsen, 2013). The species is ideal to explore how phenotypic diversity is realized within and among generations as it expresses a high degree of phenotypic plasticity and rapid intra-specific divergence, and as a result of its range and habitat use, it experiences diverse selective environments, which facilitate tracking the link between ecological processes and evolutionary outcomes (Adams, Woltering & Alexander, 2003; Garduño-Paz & Adams, 2010; Elmer, 2016; Chavarie *et al*., 2010). Given that increasing climate variability is predicted to pose a greater risk than directional climate change (Vasseur *et al*., 2014), we tested the effect of an elevated water temperature of 4°C on the short-term, plastic, phenotypic expression in an Arctic charr population that displays continuous variation. The induced temperature variation in this study is comparable in magnitude of year on year of extreme temperature fluctuations predicted by climate change models that Arctic charr populations are likely to experience (an interannual 4 ºC temperature difference is within the predicted range for the near future in northern areas; Woelders *et al*., 2018; Alley *et al*., 2003; Heino, Virkkala & Toivonen, 2009). We predict that Arctic charr individuals reared in warmer water will express greater phenotypic variation than fish at a control temperature (see Figure 1C) but that they will not display a significant shift in the mean phenotype expressed due to the short-term exposure to elevated temperatures (Figure 1B and D) (O’Dea *et al*., 2019). Overall, we aimed to: 1) quantify the among-individual morphological variation of the head of a unimodal Arctic charr population exposed to different temperature regimes during early development, 2) determine the allometry trajectories that result in any differences in expressed phenotype as evidence of differing developmental pathways in groups exposed to different temperature and 3) examine if at different temperature treatments, head shape among individuals is more analogous to the expressed phenotypes in plankton feeding (the parental phenotype) or macro-benthos feeding ecotypes in the wild.

**Figure 1.**
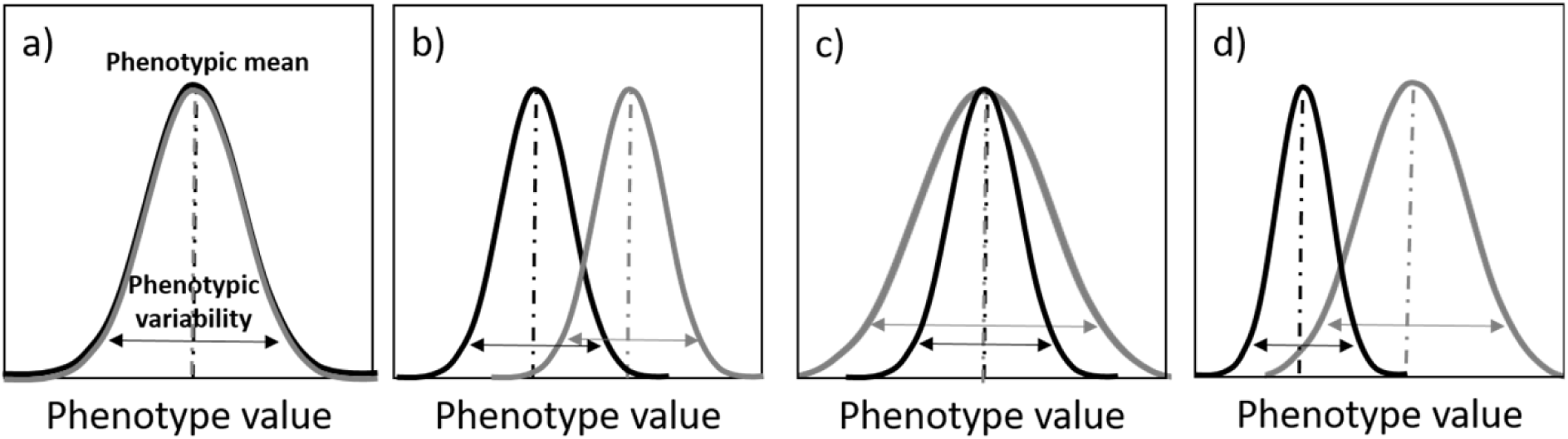
Hypothetical expressed phenotype frequency distributions shown as reaction norms resulting from an environmental change, such as temperature. Black lines represent expressed phenotypes in a population subjected to exisiting conditions and grey line represent population expression of phenotype subjected to the new environmental conditions. In panel a), the population doesn’t express any change in phenotypic mean or in phenotypic variation when exposed to the new environmental conditions. In panel b), the population shows a shift in the expressed phenotypic mean but not in phenotypic variation when exposed to the new environmental regime. In panel c), population the phenotypic mean stays the same after the environmental condition changes, but the variation in expressed phenotypic increases in the population exposed to the new environmental regime. In panel d), the population (ecotype) subjected to the new environmental conditions shift both in phenotype mean and in phenotypic variation. Direction and magnitude of variation in panel b-d are not absolute, and will be species and location dependent as well of the strength and time of exposure to the environmental variables.

## MATERIALS and METHODS

### Fish collection and rearing

Eggs from nine separate female Arctic charr were each fertilised by a single male to create nine full sibling crosses, from a morphologically unimodal, plankton feeding population that inhabits Loch Clair, Scotland. Fish were caught during spawning time from the in-flowing River Coulin (57°32.648’N, 5°19.125’W). Fertilised eggs were water hardened and transferred to incubation facilities at the Scottish Centre for Ecology and the Natural Environment, Loch Lomond. Eggs were acclimatised over a two-hour period to the new water supply and placed in mesh baskets suspended in a holding tank in a constant temperature room maintaining a water temperature of 4 °C (± 0.5 °C). Fertilisation success was greater than 95% for all family groups.

### Experimental procedure

To control for differences in development due to different temperatures, the number of degree-days (dd) were used as a measure of developmental rate. Degree-days are the cumulative count of the water temperature for a known period of time in days. Embryos reached the eyed stage after 212dd at which point they were raised to a temperature of 6 °C (± 0.5 °C) for a further 89dd, before being separated into two temperature treatment groups (n=480 per group), an ambient and an elevated temperature treatment. Developing embryos were held in equal numbers in eight replicate tanks per treatment group (n=60 per tank). Water temperatures were then either lowered by 2 °C to 4 °C or raised by 2 °C to 8 °C (± 0.5 °C) respectively. Embryos exposed to the ambient temperature (4 °C) began hatching after 367dd, the hatching period lasted 73dd (total developmental time to 100% hatch 440dd). Those exposed to the elevated temperature (8 °C) began hatching after 388dd, the hatching period lasted 50dd (total incubation time to 100% hatch 438dd). There was no significant difference in hatching rate between temperatures or replicates. When hatching was complete, temperatures were raised by 3 °C; the ambient temperature treatment to 7 °C and the elevated temperature treatment to 11 °C (± 0.5 °C). Fish became partially dependant on exogenous food at 505dd for the ambient temperature treatment and 504dd for the elevated temperature treatment. When the yolk sack was fully exhausted, 689dd for the ambient temperature treatment and 686dd for the elevated temperature treatment, temperatures were raised a further 2 °C for both treatments to 9 °C (ambient temperature treatment) and 13 °C (elevated temperature treatment). Both temperature treatments remained within ± 0.5 °C. Fish were fed four times a day to satiation at three hour intervals (± 0.5 hours) using a standard commercial 3mm hatchery sinking pellet.

### Data collection

Adult Arctic charr used as brood stock were killed using a Schedule 1 method (UK Home Office Licence Number PPL 70/8794), photographed in a lateral position on the left side before spawning. Lateral view photographs of juveniles killed using a Schedule 1 method and under licence were taken at 700dd (N=130), 1000dd (N=80) and 1400dd (N=60) using a Cannon EOS 350D digital camera, for geometric morphometric analysis. Nine consistently identifiable landmarks on the head (Figure A1) were digitised in two dimensions on each fish image using TPSdig2 (Rohlf, 2006a) and TPSutil (Rohlf, 2006b).

### Data analysis

Prior to geometric morphometric analysis, landmark data were subject to a Procrustes superimposition using MorphoJ (Klingenberg, 2011) to remove variation in the data created by size, position and orientation (Mitteroecker, 2009). The mean shape configuration was then computed and the variation around this mean calculated (Dryden & Mardia, 1998).

Following this, a single, pooled within-group regression of Procrustes co-ordinates on log centroid size was conducted in MorphoJ for samples collected at 700dd, 1000dd and 1400dd’s. The residuals from this regression provide a measure of shape free from allometric scaling (Klingenberg & McIntyre, 1998) associated with early ontogeny. The residuals from this regression were subsequently used for all further morphometric analysis.

A single Discriminant Function Analysis (1000 permutations) was conducted in MorphoJ to compare geometric morphometric data from three developmental stages (700dd, 1000dd and 1400dd) to test if the magnitude of shape difference between groups changed over time. Procrustes distance and Mahalanobis distance were used as pairwise measures of the magnitude of shape difference. Within each temperature treatment (ambient and warm), the polynomial relationship between discriminant function scores and size (represented by centroid size) were used to illustrate allometric trajectories. To decide whether allometric trajectories between temperature treatments were parallel, convergent, divergent, or common (see Figure A2), the slopes of the regressions were examined

Using the scores generated in the Discriminant Function Analysis as a measure of shape, a generalised linear mixed effect model, fitted by maximum likelihood (using the software R 3.1 for Windows (Team, 2014) and the package lme4) was used to describe the effect of temperature, exposure time (the number of dd at each developmental stage), and fish size (measured as centroid size) with replicate (tank) as a random effect, on the expression of shape. Models were simplified by removing the highest order, least significant terms, sequentially (Crawley, 2012). Terms that were removed from the model without significantly increasing model deviance using likelihood ratio tests (χ^2^) were discarded. *Post hoc* pairwise comparisons using T-test assessed if centroid size (related to body size) differed between ambient and elevated treatments at each sampling time (700, 1000, and 1400dd).

A Canonical Variate Analysis (1000 permutations) executed in MorphoJ was used to establish which group had been affected by the temperature treatment by comparing the head shape of individuals raised in ambient or elevated temperature conditions in the laboratory to their parents.

A variance ratio test was conducted using the software R 3.1 for Windows on all raw Procrustes coordinates and used to compare within-group phenotypic variation between groups.

To establish which temperature regime had stimulated expression of head shape closest to a typical plankton and macro-benthos feeding specialists from the wild, head shape of juveniles at 1400dd was compared against the head shape of two known ecologically divergent sympatric polymorphic groups of Arctic charr that show morphological and dietary specialisations on either plankton or macro-benthos feeding resources, one from Loch Rannoch (Perthshire, Scotland) (Adams *et al*., 1998) and one from Loch Dughaill (Strathcarron, Scotland) (Hooker *et al*., 2016a) using a Canonical Variate Analysis (1000 permutations) in MorphoJ. The same nine consistently identifiable landmarks used for analysing laboratory raised individuals were digitised in two dimensions on existing images of fish from Loch Rannoch and Loch Dughaill.

## RESULTS

The Discriminant Function Analysis (DFA) showed that head shape differed across all sampling periods between ambient and elevated temperature exposed fish (Procrustes distance = 0.028; P<0.01). The magnitude of shape differences however increased over time (at 700dd, Procrustes distance = 0.036; P<0.01; at 1000dd, Procrustes = 0.040; P<0.01; at 1400dd, Procrustes distance = 0.061; P<0.01; Figure 2).

**Figure 2.**
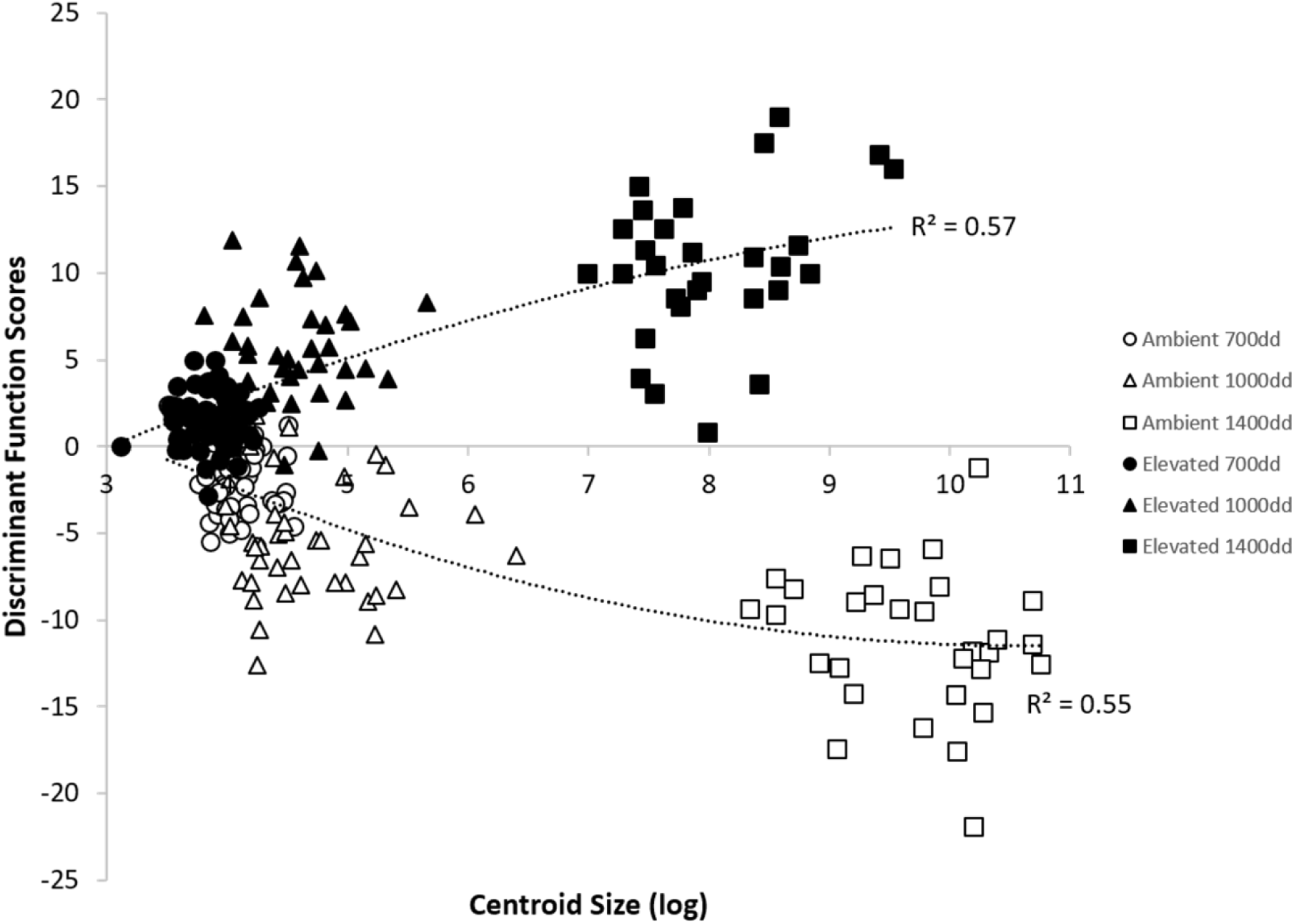
Individual Discriminant Function Scores (DF1) for head shape of Arctic charr of different sizes (represented by centroid size). Three sample periods are given, 700dd are represented by circles, 1000dd by triangles and 1400dd by squares. Ambient exposed fish are denoted with open symbols; elevated temperature fish are denoted with closed symbols. Polynomial regressions for both temperature treatments are showing allometric trajectories toward phenotypic divergence of head shape changes (see Figure A2).

Discriminant Function Analysis of shape only, correctly assigned 73.8% of ambient temperature fish and 82.8% of elevated temperature fish (from all sampling periods combined) to the correct temperature exposure. In general, fish exposed to elevated temperature had a more rounded head and sub-terminal mouth than ambient temperature fish (Figure A3). Individuals in both temperature treatments displayed allometric trajectories with polynomial relationships of head shape on size differing from 0 (P<0.05); centroid size explained 55% of variation in Discriminant Function Score of head shape in the ambient temperature treatment group (R^2^ =0.55) and 57% in the elevated temperature treatment group (R^2^=0.57). Regression slopes indicated that allometric trajectories were divergent (Figure 2 and Figure A2).

Using discriminant function scores as a measure of shape in a mixed model, temperature had a highly significant effect on head shape (likelihood ratio ***χ*2** = 39.81, p<0.01) (Table 1). There was a significant interaction between Exposure time and Centroid Size (likelihood ratio ***χ*2** = 7.23, p = 0.01) (Table 1). The interaction between exposure (number of degree-days) and fish size (centroid size) was negative. Thus, the difference in morphology between fish raised on each temperature regime was greater in larger fish, but the rate of divergence decreased as fish became larger; indicating that speed of divergence caused by different temperature regimes is greater during early ontogenetic stages. Pairwise comparisons of centroid size between temperature treatments at each sampling time, showed a difference in size at 700dd (t = 1.98, p < 0.01) and at 1400dd (t = 2.0, p < 0.01), but no size differences were found at 1000dd (P < 0.05). At 700dd and 1400dd, individuals raised at ambient temperature were larger than individuals exposed to elevated temperature.

**Table 1.** Parameter estimates from the minimum adequate generalized linear mixed model describing the effect of temperature, exposure and centroid size on shape DF scores).

| | Estimate | SE | $X^2$ | $P$ |
| --- | --- | --- | --- | --- |
| Intercept (Ambient) | -6.84 | 1.91 | - | - |
| Exposure time | 0.002 | 0.002 | - | - |
| Elevated temperature | 2.36 | 0.21 | 39.81 | <0.01 |
| Centroid size | 1.63 | 0.48 | - | - |
| Exposure time : Centroid Size | -0.0012 | 0.0005 | 7.23 | <0.01 |

The variance ratio test found the phenotypic variation within treatment groups was higher in ambient temperature fish compared with elevated temperature fish (p = <0.01). For ambient temperature, variance in Procrustes scores (SS distance/(n-1)) was 0.0046 at 700dd, 0.0032 at 1000dd, and 0.0018 at 1400dd. For elevated temperature, variance of Procrustes scores (SS distance/(n-1)) was 0.0038 at 700dd, 0.0039 at 1000dd, and 0.0016 at 1400dd.

Canonical Variate Analysis (CVA) found that the head shape of fish exposed to the ambient temperature treatment was not significantly different to that of their parents (Procrustes distance = 0.014; *P*=0.38), however elevated temperature fish were significantly different (Figure A4; Procrustes distance = 0.044; *P=*<0.001).

Using a CVA to compare experimental (Coulin) charr from both temperature groups to benthic and pelagic foraging specialists from Loch Rannoch and Loch Dughaill, we found that ambient temperature exposed fish expressed a phenotype closer to the pelagic ecotype and fish exposed to elevated temperature displayed a phenotype closer to benthic ecotype for both comparisons (Figure 3 and Table 2). CV1 mostly captures shape differences between fish of different origin, experimental charr and Loch Rannoch charr (Figure 3A) and experimental and Loch Dughaill charr (Figure 3B). CV2 however captures within group shape differences between fish raised at ambient or elevated temperatures, benthic and pelagic ecotypes from Loch Rannoch (Figure 3A) and benthic and pelagic ecotypes Loch Dughaill (Figure 3B).

**Table 2.**
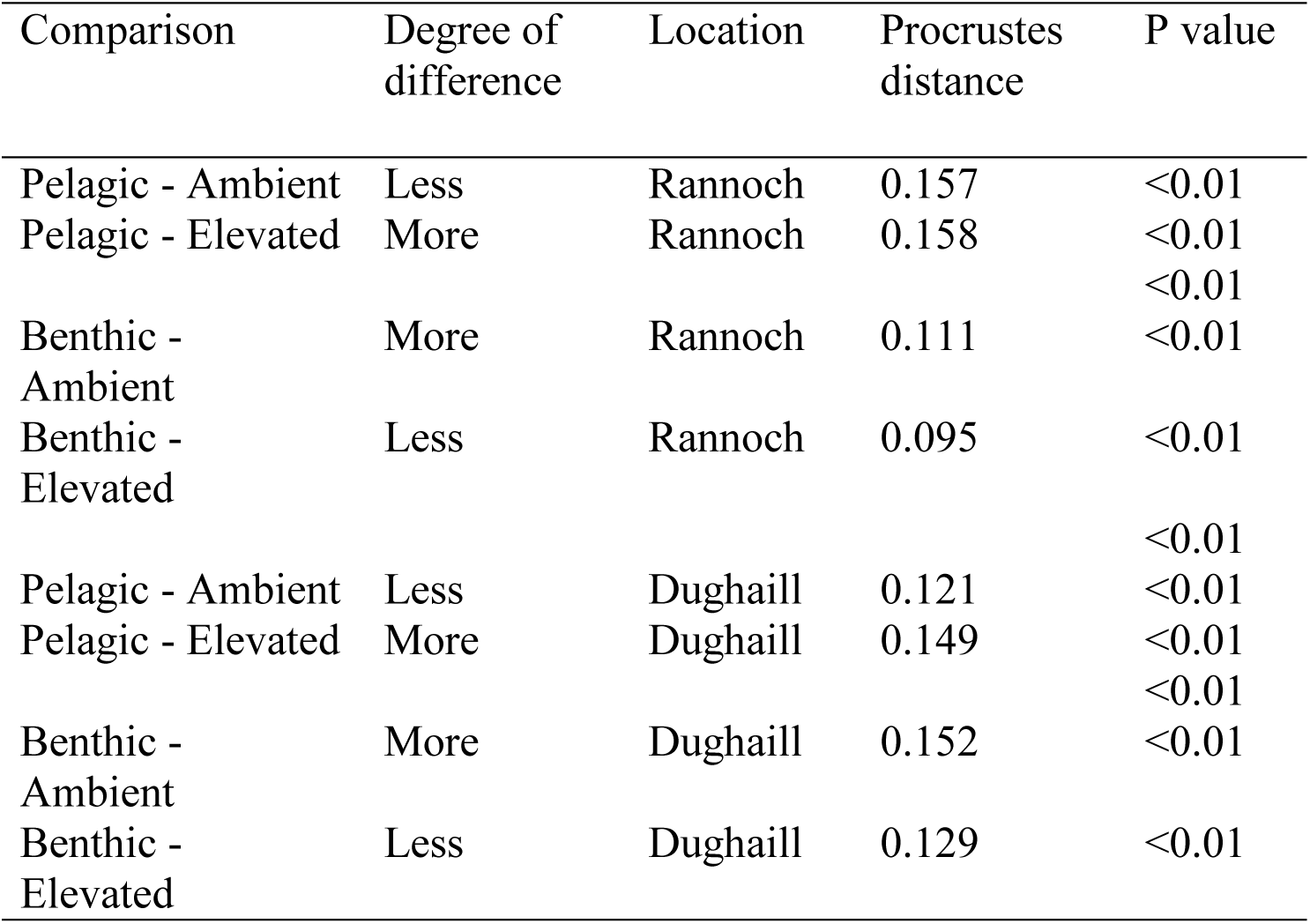
Results from the CVA comparing ambient and elevated temperature raised fish to wild benthic and pelagic ecotypes from Loch Rannoch and Loch Dughaill.

**Figure 3.**
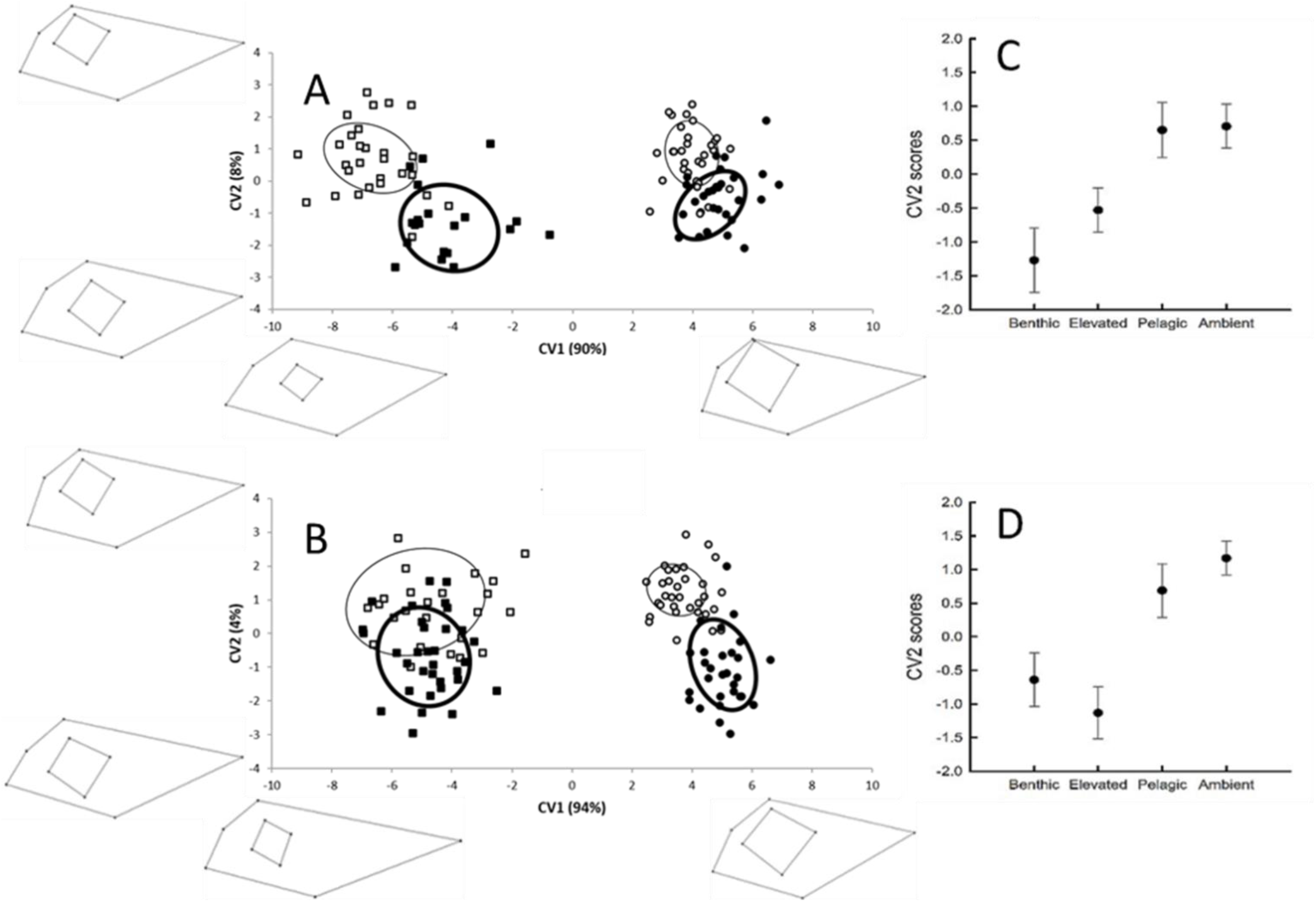
Canonical Variate Analysis of 1400dd ambient and elevated temperature exposed Arctic charr and sympatric polymorphic populations of Arctic charr from; A) Loch Rannoch, B) Loch Dughaill. Elipses represent 95% confidence limits for means. Polymorphic charr populations are represented by squares and experimental Arctic charr by circles. Open symbols denote wild plankton feeding Arctic charr and Arctic charr raised at an ambient temperature. Closed symbols denote benthic feeding Arctic charr and Arctic charr raised at an elevated temperature. X and Y axis percentages denotes the amount of variation for each canonical variate. Wire frames on CV1 are scaled at −8 and +8, wire frames for CV2 are scaled at −4 and +4. C) and D) show mean (+/- 95% CL) for CV2 scores.

## DISCUSSION

Variation in climate can occur at different temporal scales (e.g. diurnally, intra-seasonally or inter-annually), and extreme temperature anomalies are now occurring more regularly and are predicted to increase in frequency in the future (Scheepens, Deng & Bossdorf, 2018). Accordingly, in this study we demonstrate clear differences of an environmentally contingent expressed phenotype in an ectotherm subjected to a severe climate event. Our results demonstrate a strong and immediate influence of temperature on the direction, magnitude, and variation in the expression of phenotypic plasticity in a cold-water specialist species where the population in the wild displays continuous variation (i.e., unimodal phenotype; Klemetsen, 2010; Klemetsen, 2013). Elsewhere studies on plasticity during development have shown responses to alternative environmental inputs, resulting in dramatic morphological differences among individuals (Morris, 2014; West-Eberhard, 2003). Here, offspring of wild Arctic charr raised at elevated temperatures expressed a mean phenotype different to their siblings raised at an ambient temperature but also to that of their parents. The magnitude of this morphological response mimicking the level of divergence observed in wild Arctic charr populations that express sympatric foraging specialist ecotypes showing a polymodal phenotype distribution (Skulason & Smith, 1995). When compared to polymorphic populations that show distinct and stable foraging specialisms, individuals raised in the elevated temperature treatment expressed a phenotype analogous to a benthivorous ecotype rather than that of their parental pelagic form (Adams *et al*., 1998; Hooker *et al*., 2016b). The head of fish exposed to elevated temperature was shorter, more robust, with a rounder snout; features that are typical of littoral macro-benthos feeding specialists (Figure A3) in several post-glacial fishes (Robinson & Parsons, 2002; Bolnick & Lau, 2008). In contrast, ambient temperature exposed fish expressed an elongated and pointed head phenotype, which is more typical of fish showing pelagic planktivory in the wild (Schluter, 2000; Taylor, 1999).

Exposure to different environmental conditions is known to have asymmetrical impacts on the stability of developmental pathways during early life-stages (Robinson & Wardrop, 2002; Svanbäck *et al*., 2009; Lazić *et al*., 2013). Considering that temperature is one of the most widely reported environmental factors driving plastic responses in fish (Pittman *et al*., 2013), Arctic charr individuals subjected to higher temperature developed at a faster rate (Campbell, 2019). Developmental stability is known to limit the phenotypic expression from a single genotype, thus it can modulate the magnitude and structure of phenotypic variation among individuals and cohorts (Lazić *et al*., 2015; Willmore *et al*., 2006). In some cases, the expressed phenotype of an organism can be significantly altered by a single short-term environmental perturbation of the timing or rate during development; such events are thought to be a route to rapid phenotypic novelty, the raw material for selection (Parsons *et al*., 2011). Several mechanisms, such as heterochrony, are known to drive the plastic expression of phenotypes through the effects of temperature exposure on developmental pathways (Georga & Koumoundouros, 2010; Angilletta, Steury & Sears, 2004; Ramler *et al*., 2014). Heterochrony can disrupt the onset and termination of developmental processes and the rate at which these processes occur. Developmental pace in structurally important features, such as bone metabolism, are fundamentally linked to the form of morphological plasticity examined in this experiment (Campinho, Moutou & Power, 2004; Sfakianakis *et al*., 2004). Therefore, temperature effects mediated through heterochrony has the potential to result in skeletal development of different craniofacial features in Arctic charr (Parsons & Albertson, 2009; Campbell, 2019).

Early development represents a critical life-stage for the emergence of phenotypic variation. During ontogeny, the environment is likely to have a greater influence on a range of traits and the magnitude of the variation of expression in a population (Gillooly *et al*., 2001; Ackerly & Ward, 2016; Imsland *et al*., 2005; Parsons, Skúlason & Ferguson, 2010). The thermal history experienced by an individual during early development is known to be a key factor defining adult phenotype (Georga & Koumoundouros, 2010; Angilletta, Steury & Sears, 2004; Ramler *et al*., 2014). It is particularly during early development that environmental signals can influence developmental processes through epigenetic mechanisms, influencing the expressed phenotype (Duclos, Hendrikse & Jamniczky, 2019). This observed effect on the phenotype in this study was cumulative, with divergence between treatment groups increasing over time. However, the rate of divergence decreased with time, suggesting that the effect of temperature on the expression of phenotypic differences occurred predominant at early life stages. This indicates that the timing of exposure to different environmental conditions by individuals is important in the initiation of a plastic response and its magnitude.

Temperature exposure in early life is known to result in epigenetic effects on traits such as growth, with consequences that can range from short-term to transgenerational (Burton & Metcalfe, 2014). Our results suggest that temperature-induced variation in developmental pathways results in divergent plastic allometric trajectories in the early development of Arctic charr. Allometry, can been seen as a canalising process and an interacting agent that can steer morphological variation (Franklin *et al*., 2018; Simonsen *et al*., 2017). However, the opposite is also true, whereas allometric trajectories can deviate from the “usual” shape-size relationship (e.g., whereas size increases, shape changes under a fixed allometric relationship), deviation from this fixed allometry relationship could serve as additional measure of development instability (Lazić *et al*., 2015). Consequently, growth trajectories manifested as allometric scaling provide the potential to express different phenotypes, which increase the potential for novelty (Frankino *et al*., 2005). Altogether, both heterochrony and allometry are involved in determining the degree of phenotypic variation generated during growth and morphological development. In this study, the expression of phenotypic variation observed appears to be influenced by temperature with concomitant developmental effects (Lazić *et al*., 2015; Westneat, Wright & Dingemanse, 2015).

The potential of Arctic charr to display phenotypic variation is considerable, whether expressed variation is continuous (e.g., unimodal) or discontinuous (e.g., polymodal) (Klemetsen, 2013). Populations that express high level of variability in natural conditions show higher means and greater variance in the plastic expression of phenotype in response to different temperatures (O’Dea *et al*., 2019; Reed *et al*., 2010). Although our study also showed a significant plastic response resulting in a shift in the mean phenotype expressed, there was also a decrease in variation of the phenotypes expressed among individuals raised at higher temperature treatment, which contrasts with previous work. Increased phenotypic variance in response to elevated temperatures has been suggested to be the reflection of a reduction in genotype precision (e.g., canalization and developmental stability), this lack of precision resulting in a spreading of reaction norms (O’Dea *et al*., 2019; Campbell, 2019). Our opposing result could be the result of a non-linear reaction norm of head shape on size. In this scenario a specific range of temperature exposures (either colder or warmer) has a different magnitude effect on phenotypic mean and variation (see description of how this may occur in Figure 1 in Ramler *et al*., 2014). Another, not mutually exclusive possibility, is that despite both treatments being sampled after the same number of degree-days (a proxy for growth opportunity; Neuheimer & Taggart, 2007) had elapsed, individual responses in the two temperature groups was partly the result of a size effect. For an unknown reason, in our experiment, growth opportunity and the actual realized body growth were uncoupled. Contrary to the predictions of Neuheimer and Taggart (2007), the effect of time and temperature are thus not necessary equal in the promotion of body size and anatomical structures (e.g., heterochrony and allometry). Consequently, if variance is cumulative with the increase of size (Gillooly *et al*., 2002), this could partly explain the discrepancies between our predictions and findings because ambient temperature treatment individuals were generally larger by the end of the experiment.

### Broader implications

Environmentally mediated variation in phenotypic expression is a fundamental attribute of many organisms, which can result in both short and long-term ecological evolutionary consequences for species that display plasticity. The disjunctive nature of the freshwater environment that lacustrine Arctic charr occupy means that, in the absence of migratory routes for range adjustment, this species will particularly need to cope to abrupt changing novel conditions (O’Dea *et al*., 2016; Collins *et al*., 2013). High levels of plasticity is one way in which they might do this. Yet, the role of phenotypic plasticity as a potential mitigator of climate change is contentious, with both positive and negative effects possible. We demonstrate that the expressed phenotypic mean in Arctic charr can respond to an environmental cue rapidly and with a magnitude that is highly likely to have functional consequences (i.e. temperature-mediated shift from a pelagic-like towards a benthic-like phenotype). In the context that even subtle differences in individual’s morphology can influence feeding efficiency and specialization (Garduño-Paz & Adams, 2010), the potential of a fitness-buffering effect of plasticity can easily fluctuate, depending if environmental cues and selective filters become decoupled or not (Reed *et al*., 2010; O’Dea *et al*., 2016). Consequently, important questions arise when extrapolating our results into the wild. Would such plastic phenotypic change result in an “overshoot” of the optimum phenotype (e.g., would such processes produce phenotype-environment mismatches) or would it buffer against rapid fluctuations in the environment?

While a plasticity response occurs within a single cohort, another important question at hand is what would be the consequences to subsequent generations (e.g., transgenerational plasticity effects)? The high magnitude of the short temporal environmental oscillations, for instance extreme environmental events, can induce a greater set of phenotypic responses than a small or slower change (e.g., our tested 4 ºC variation scenario; Donelson *et al*., 2018). More gradual or stepwise change across generations have the potential to produce different phenotypic results compared to a single large change within a cohort (e.g., reproductive capacity of individuals subjected to a 3ºC elevated temperature within a generation compared with a 1.5ºC elevation over two generations; Donelson *et al*., 2016). In reality the probability of occurrence of alternative effects of phenotypic plasticity (positive or negative) on responses to temperature change is likely to be an interaction between the capacity of the organism for phenotypic plasticity (both within-generation and transgenerational plasticity), the speed of mean change in the environment (e.g., temperature), and the degree of short interval variation in the environment (Reed *et al*., 2010; Donelson *et al*., 2018).

The last component emerging from this experiment with evolutionary consequences for a population in the face of changing selection regimes by affecting population dynamics is the phenotypic variance (Chevin, Lande & Mace, 2010; Reed *et al*., 2010). In this study, we showed a decrease in variation in expressed phenotype at elevated temperature. The decrease in the variation in the phenotypes expressed among individuals raised in the elevated temperature treatment could reduce the range of phenotypes for selection to act upon if this effect occurs in the wild. Accordingly, phenotypic variance has the potential to be a very important component providing a population with scope for recovery from rapid (possibly transient) environmental change. Consequently, a reduction or an increase of expressed phenotypic variance under climate change scenarios has important consequences at the population level should not be underestimated.

## CONCLUSION

The ecological and evolutionary consequence of rising temperature on Arctic charr will depend on many factors (e.g., population specific lethal limits or growth optima, interspecific dynamics, and changing disease risk; Crozier & Hutchings, 2014). Here, we demonstrate the capacity of Arctic charr for a rapid plastic response to the effect of a severe temperature fluctuation in effect over periods of less than a single year. The parental phenotype mean was shifted within a generation, but also individuals showed a marked decline in in the expression of intra-populaton phenotypic variation. Such variation has been recently shown to have multi-generational effects on survival in another ectotherm the minnow *Phoxinus phoxinus* (Raffard *et al*., 2019). Temperature varies in a stochastic manner resulting in year-to year variation around a coarse-scale temporal trend. Inter-annual temperature variation can modify a cohort’s phenotype in a population, consolidating the role of plasticity in species resilience to rapid environmental change, as genetic changes are not expected to be capable of responding over such short time scales (Merilä & Hendry, 2014).

Phenotypic plasticity is thus likely to be particularly important for population persistence in new and changing environments and where the pace of change is relatively rapid (Morris, 2014). Contemporary temperature-driven divergence in Nordic freshwater fishes have been reported (Kavanagh *et al*., 2010). Whether this flexible response will be common within and among cold-water species is unknown. Nonetheless, the fact remains that the majority of landlocked Arctic charr populations are currently planktivorous (Maitland & Adams, 2018). By confirming the potential of a rapid and marked change in morphology with higher temperature, we provide evidence that a substantial portion of Arctic charr populations worldwide are likely to be subjected to phenotype expression changes toward a benthic-like ecotype under climate change scenarios. A similar effect may also occur in other salmonids and other cold-water species.

## ACKNOWLEDGMENTS

The authors would like to acknowledge Travis van Leeuwen, Luc Bussierre, Alex Lyle, Jennifer Dodd, Martin Hughes, Peter Cunningham (Wester Ross Fisheries Trust) and The Coulin Estate where fish were collected.

## DATA AVAILABILITY

Upon acceptance, data will be available in dryad.

## Appendix

**Figure A1.**
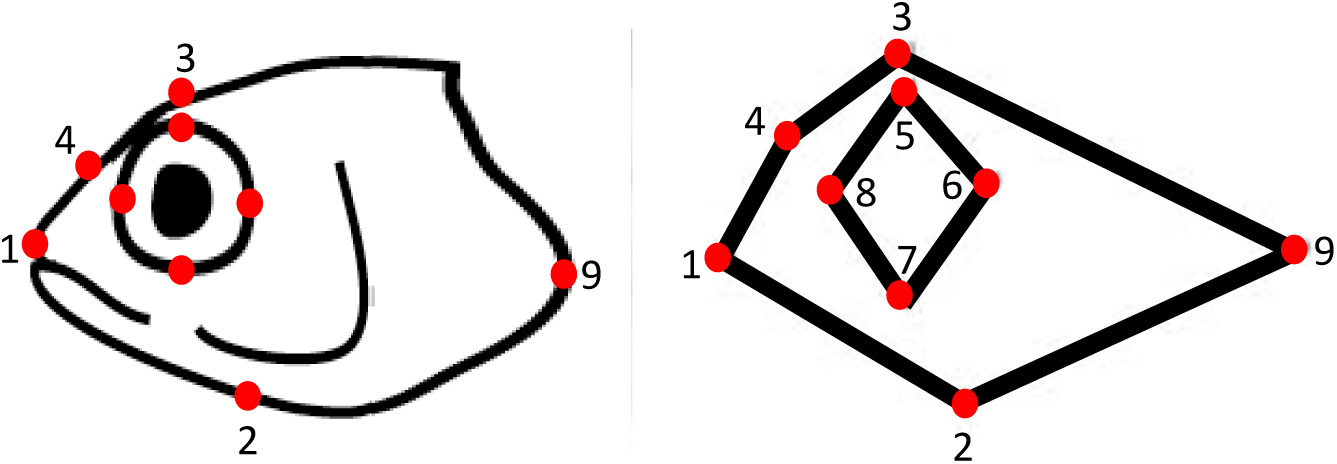
Nine landmarks were used to characterise the shape of the head; Landmark (LM) 1, the tip of the nose; LM2, the most posterior part of the upper jaw; LM3, edge of cranium directly above the eye; LM4, edge of cranium at the central point between LM3 and LM1 at a 90 degree angle; LM5-8, most upper, posterior, lower and anterior parts of eye respectively; LM9, most posterior part of the gill operculum.

**Figure A2.**
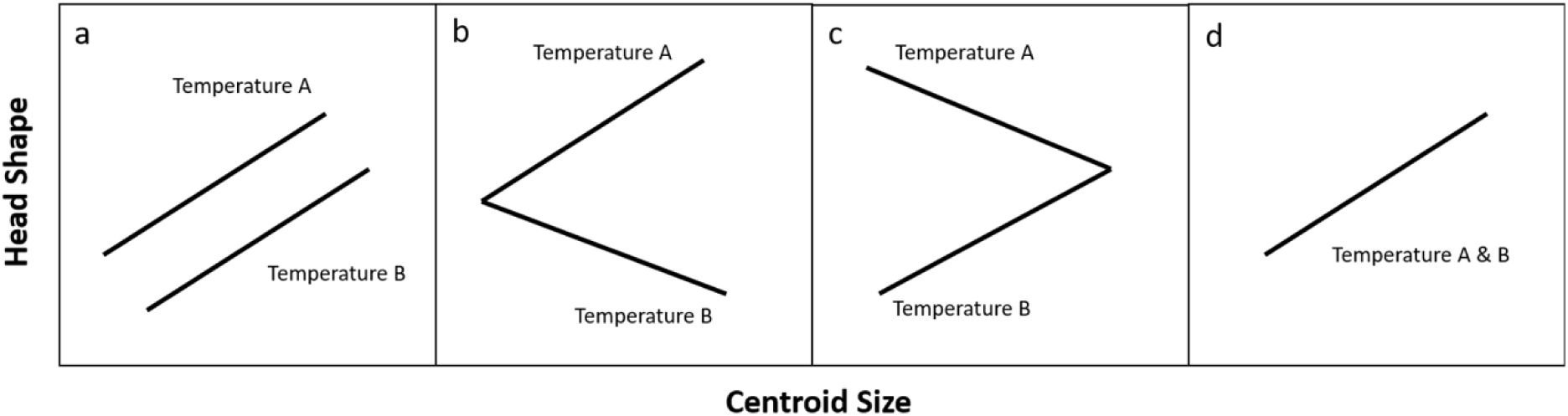
Possible pattern of allometric trajectories between temperature treatments of this study, figure is modified from Simonsen *et al*. (2017). Allometric trajectory patterns can be a) parallel, b) divergent, c) convergent, or d) common.

**Figure A3.**
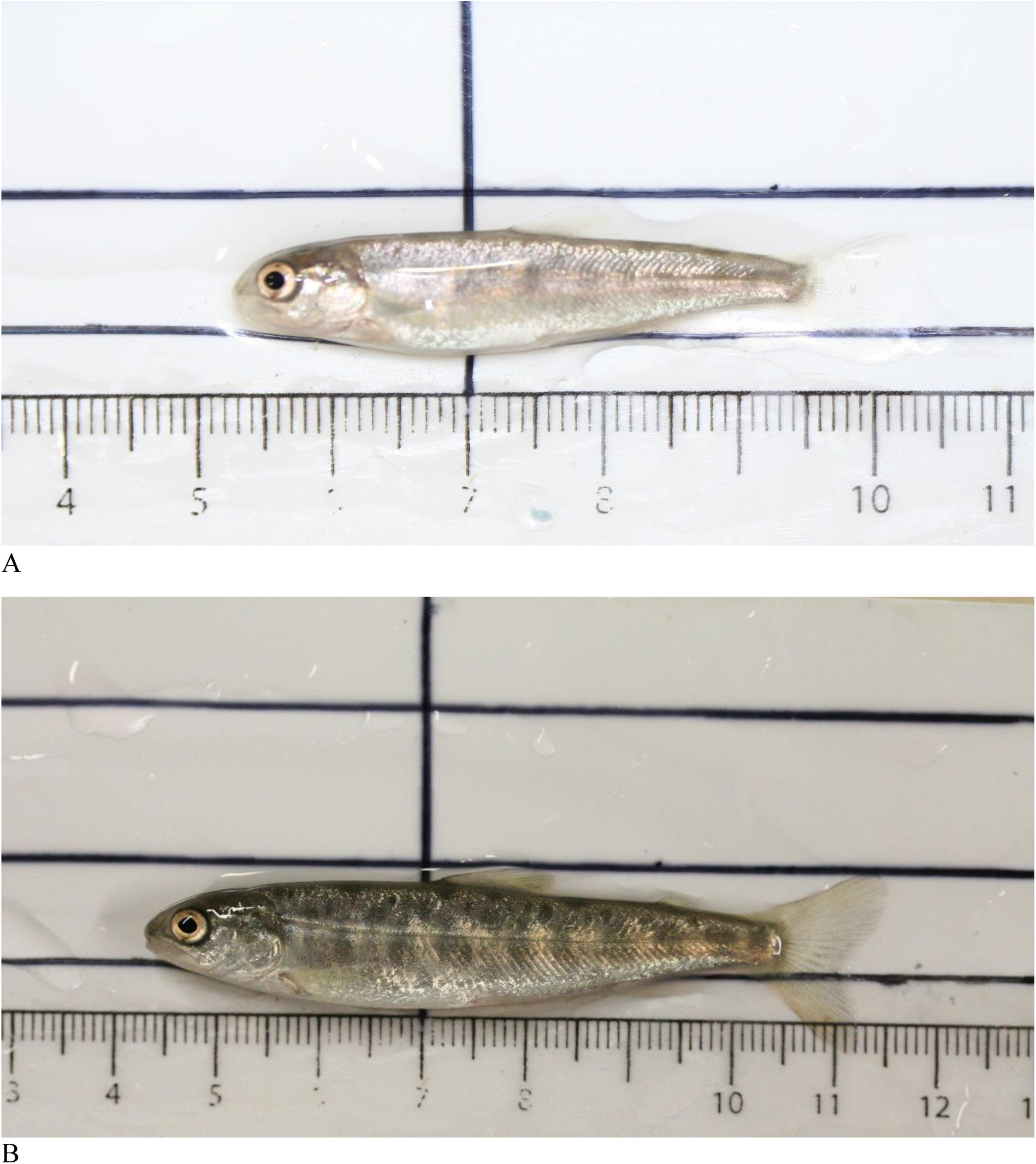
Individuals sampled at 1400dd, from the group a) raised at elevated temperature, with a morphology analogous to benthic feeding wild ecotype and b) raised at ambient temperature, with a morphology analogous to plankton feeding wild ecotype (parental population).

**Figure A4.**
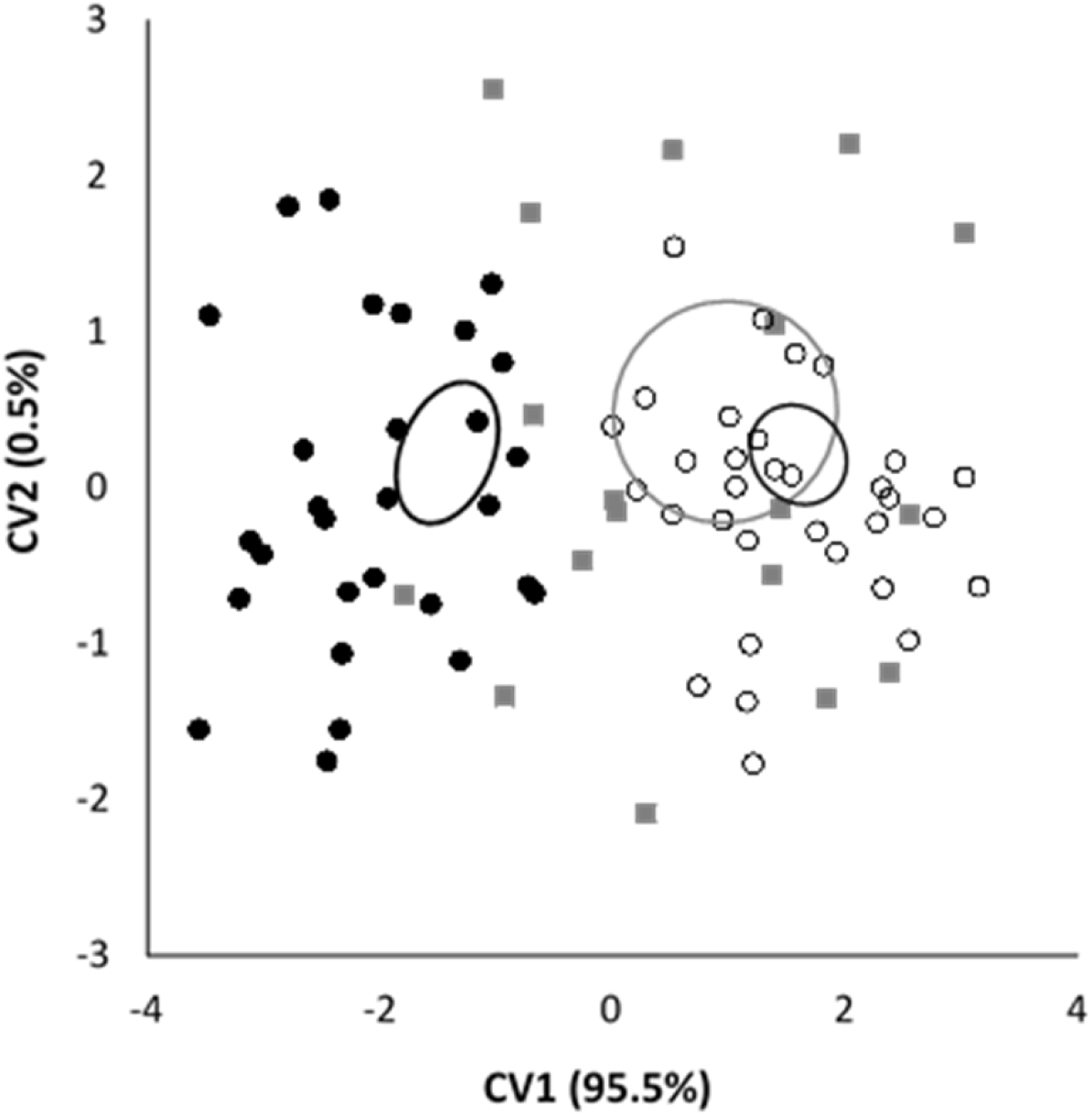
CV1 and CV2 scores derived from Canonical Variate Analysis of: parent fish (grey squares), ambient temperature exposed fish sampled at 1400dd (open circles) and elevated temperature exposed fish sampled at 1400dd (closed circles) with 95% confidence elipses of centroid. Percentage on X and Y axes denotes the amount of variation explained by each canonical variate.

